# A conserved acidic cluster motif in SERINC5 confers resistance to antagonism by HIV-1 Nef

**DOI:** 10.1101/590646

**Authors:** Charlotte A. Stoneham, Peter W. Ramirez, Rajendra Singh, Marissa Suarez, Andrew Debray, Christopher Lim, Xiaofei Jia, Yong Xiong, John Guatelli

## Abstract

The cellular protein SERINC5 inhibits the infectivity of diverse retroviruses and is counteracted by the glycoGag protein of MLV, the S2 protein of EIAV, and the Nef protein of HIV-1. Determining regions within SERINC5 that provide restrictive activity or Nef-sensitivity should inform mechanistic models of the SERINC5/HIV-1 relationship. Here, we report that deletion of the highly conserved sequence EDTEE, which is located within a cytoplasmic loop of SERINC5 and is reminiscent of an acidic cluster membrane trafficking signal, increases the sensitivity of SERINC5 to antagonism by Nef while having no effect on the intrinsic activity of the protein as an inhibitor of infectivity. The effects on infectivity correlated with enhanced removal of the ΔEDTEE mutant relative to wild type SERINC5 from the cell surface and with enhanced exclusion of the mutant protein from virions by Nef. Mutational analysis revealed that the acidic residues, but not the threonine, within the EDTEE motif are important for the relative resistance to Nef. Deletion of the EDTEE sequence did not increase the sensitivity of SERINC5 to antagonism by the glycoGag protein of MLV, suggesting that its virologic role is Nef-specific. These results are consistent with the reported mapping of the cytoplasmic loop that contains the EDTEE sequence as a general determinant of Nef-responsiveness, but they further indicate that sequences inhibitory to as well as supportive of Nef-activity reside in this region. We speculate that the EDTEE motif might have evolved to mediate resistance against retroviruses that use Nef-like proteins to antagonize SERINC5.

**Importance:** Cellular membrane proteins in the SERINC family, especially SERINC5, inhibit the infectivity of retroviral virions. This inhibition is counteracted by retroviral proteins, specifically HIV-1 Nef, MLV glycoGag, and EIAV S2. One consequence of such a host-pathogen “arms race” is compensatory change in the host antiviral protein as it evolves to escape the effects of the viral antagonist. This is often reflected in a genetic signature, positive selection, which is conspicuously missing in *SERINC5*. Here we show that despite this lack of genetic evidence, a sequence in SERINC5 nonetheless provides relative resistance to antagonism by HIV-1 Nef.

## Introduction

HIV-1 is a complex retrovirus, encoding “accessory” genes that evolved to enhance viral fitness in response to host-selective pressures (1). The accessory gene *nef* accelerates *in vivo* pathogenesis and progression to AIDS, despite being non-essential for viral propogation in cell-culture (2–4). Expression of the Nef protein occurs early during the viral replication cycle, preceding the expression of structural proteins such as the envelope glycoprotein (Env) and preceding virion assembly (5). Post-translational myristoylation on an N-terminal glycine residue enables Nef to associate with lipid membranes (6), where it modulates the trafficking of host proteins to promote immune evasion. Nef-activities include down-regulation of the HIV receptor CD4 (6) and the major histocompatibility complex I (MHC-I) from the cell surface (7). To modulate CD4, Nef uses a di-leucine-based motif to recruit components of the cellular protein sorting machinery, specifically the clathrin-Adaptor-Protein complex 2 (AP-2), to induce the endocytosis of CD4 and ultimately target it to the multivesicular body (MVB) pathway for lysosomal degradation (8–11). CD4 modulation is also an activity of the HIV accessory protein Vpu; together the activities of Vpu and Nef prevent CD4 and Env from interacting in the virion-producer cell. This ensures the proper maturation of Env, preventing CD4 from inhibiting virion-infectivity and from triggering the exposure of CD4-dependent epitopes in Env that are good targets for host humoral immunity (12–17). In contrast to the above consensus regarding CD4, two mechanisms have been proposed for Nef-mediated modulation of MHC-I: 1) Nef utilizes the clathrin adaptor AP-1 to bind newly synthesized and antigen-loaded MHC-I molecules within the *trans*-Golgi network (TGN) to target them for eventual lysosomal degradation (18); and 2) Nef accelerates the internalization of MHC-I from the cell-surface via a PI3-kinase-regulated and ARF6-mediated pathway to promote sequestration of MHC-I within the TGN (19). Either of these mechanisms could lead to a reduction of MHC-I molecules at the cell surface and resistance of HIV-1 infected cells to killing by cytotoxic T lymphocytes (20).

Another highly conserved activity of Nef is the enhancement of virion-infectivity (21, 22). This activity is preserved among *nef* alleles obtained from HIV-1-infected individuals at different stages of disease progression, suggesting that it is important both for transmission and for persistent infection (23). The infectivity-effect is dependent on specific regions within Nef, all of which are also required for the modulation of CD4, including the above noted di-leucine-based motif. Components of the cellular endocytic machinery (AP-2, Dynamin 2, and clathrin) are also required (24, 25). Nef must be expressed within virion-producer cells to enhance infectivity; its presence in target cells and in virions is dispensable (26, 27). These observations led to the hypothesis that Nef prevents a cell surface “infectivity-inhibiting-factor” from incorporating into virions.

Serine incorporator 3 and 5 (SERINC3 and SERINC5) were identified as such factors; they are transmembrane proteins that incorporate into virions and potently inhibit the infectivity of retroviruses (28, 29). Nef counteracts this by removing SERINC3 and SERINC5 from the plasma membrane in an AP-2 dependent manner (29). Nef’s ability to enhance infectivity depends quantitatively on the relative sensitivity or resistance of the tested Env protein to inhibition by the SERINCs: the Nef-effect is greatest when the matching Env protein is sensitive to SERINC5 (28, 29). This sensitivity in turn appears to correlate directly with the “openness” of the Env trimer and consequently with the sensitivity of the Env to neutralizing antibodies that are selectively active against more “open” trimers (30).

SERINC3 and SERINC5 are members of a conserved family of proteins whose cellular function includes phospholipid biosynthesis, specifically the incorporation of serine into membrane lipids (31). Nonetheless, SERINC5 does not appear to alter the lipid composition of virions (32). Instead, SERINC5 inhibits the fusion of virions with target cells, potentially by functionally inactivating sensitive Env trimers (33). Nef prevents the incorporation of SERINC5 into virions, presumably by physically interacting with SERINC5, and then stimulating its endocytosis and sending the protein toward lysosomal degradation (34). SERINC5 antagonists have been identified in retroviruses other than HIV and SIV. These include the glycoGag protein of Murine Leukemia Virus (MLV) (35) and the S2 protein of Equine Infectious Amenia Virus (EIAV) (36). While Nef and glycoGag are structurally unrelated, the mechanisms by which they counteract SERINC5 seem similar: endocystosis and lysosomal degradation (37). Despite this scenario of host-pathogen conflict between the SERINCs and retroviral proteins, *SERINC3* and *SERINC5* do not appear to be under positive selection at the protein-level, at least not to the extent observed for other anti-retroviral restriction factors such as *TRIM5α* or *BST-2* (38)

The goal of this study was to determine whether a potential membrane trafficking signal in SERINC5, reminiscent of an acidic cluster sorting motif, supported the activity of Nef. This sequence, EDTEE, is within the same cytoplasmic loop that has recently been shown to be a determinant of Nef-sensitivity (39). The hypothesis that this sequence would support Nef-activity is consistent with the roles of sequences reminiscent of sorting motifs in other Nef-targets, such as the key tyrosine in the cytoplasmic domain of the class I MHC α chain and the di-leucine motif in the cytoplasmic domain of CD4 (8, 40, 41). Paradoxically, we found that rather than supporting Nef-activity, the EDTEE sequence instead provided a degree of protection against Nef: lack of the EDTEE sequence enhanced Nef-activity as an antagonist of SERINC5. The relatively increased infectivity of virions produced in the presence of Nef and SERINC5 lacking the EDTEE sequence correlated with more efficient exclusion of SERINC5 from virions and more efficient downregulation of cell surface SERINC5 by Nef. This enhanced-response phenotype appeared to be specific to Nef; deletion of the EDTEE sequence slightly impaired rather than enhanced the activity of glycoGag as an antagonist of SERINC5. We speculate that the EDTEE region may have specifically evolved to render Nef proteins less active SERINC5-antagonists.

## Materials and Methods

### Cells

HEK293 (obtained from Dr. Saswati Chaterjee) (42) and HeLa TZM-bl cells (obtained from Dr. John Kappes via the NIH AIDS Reagent Program) were cultured in DMEM media supplemented with 10% FBS and 1% Penicillin/Streptomycin. HeLa P4.R5 cells (obtained from Dr. Ned Landau) were maintained in DMEM media supplemented with 10% FBS, 1% Penicillin/Streptomycin, and 1 µg/ml puromycin. A leukemic T cell clone (Jurkat E6.1) lacking endogenous levels of SERINC3 and 5 (termed JTAg *S3/5* KO) was a gift from Dr. Heinrich Gottlinger. These cells were cultured in complete RPMI media: 10% FBS and 1% Penicillin/Streptomycin.

### Plasmids

The proviral plasmids pNL4-3 and pNL4-3ΔNef have been described previously (21, 43, 44). The pNL4-3-derived plasmids lacking *env* (“DHIV”) or lacking both *env* and *nef* genes (“DHIVΔNef”) were gifts from Dr. Vicente Planelles (45). The plasmid pCINeo-VRE (pVRE) contains the sequence of NL4-3 from 9 bp upstream of the Rev start codon to the *Xho*I site in Nef (100 bp downstream of the Nef start codon), and encodes Vpu, Rev and Env. The empty vector pBJ5 and the pBJ5-HA-gg189 plasmid containing an HA-tagged minimal active truncated form (the N-terminal 189 residues) of MLV glycoGag were a gift from Dr. Massimo Pizzato. The plasmid pBJ5-SERINC5-iHA was a gift from Dr. Heinrich Gottlinger (28). This SERINC5 plasmid contains an HA tag located between residues 290 and 291 in extracellular loop 4 of the protein. pBJ5-SERINC5-iHA ΔEDTEE, pBJ5-SERINC5-iHA AATAA and pBJ5-SERINC5-iHA EDAEE, containing mutations within amino acids 364-368 in human SERINC5, were generated using site-directed mutagenesis (QuikChange, Agilent Technologies) using the following primers: for ΔEDTEE: CTTCAGTCCTGGTGGACAGCAGCCGGGGAAG and CTTCCCCGGCTGCTGTCC-ACCAGGACTGAAG; for AATAA: GTCCTGGTGGAGCCGCCACTGCAGCG-CAGCAGCCG and CGGCTGCTGCGCTGCAGTGGCGGCTCCACCAGGAC; for EDAEE: CTGGTGGAGAGGACGCTGAAGAGCAGCAG and CTGCTGCTCTT-CAGCGTCCTCTCCACCAG. The plasmid pcDNA3.1-SERINC5-VN-HA was a gift from Dr. Yonghui Zheng (34) and was used for the Bi-molecular Fluorescence Complementation (BiFC) assays. We used the mutagenic primers above to construct pcDNA3.1-SERINC5ΔEDTEE-VN-HA, pcDNA3.1-SERINC5 EDAEE-VN-HA and pcDNA3.1-SERINC5 AATAA-VN-HA. The plasmid pcDNA3.1-Nef_SF2_-V5-VC was a gift from Dr. Thomas Smithgall and has been described previously (46). To construct pcDNA3.1-Nef_NL43_-V5-VC or a myristoylation defective Nef (pcDNA3.1-Nef_NL43_ G2A-V5-VC), a 621 bp PCR product bearing *Not*I and *Eco*RI restriction sites was generated using template plasmids containing the NL4-3 wildtype or mutant Nef alleles (pCI-NL) (24). The sense PCR primer for wildtype was AGATTCGCGGCCGC-ACCATGGGTGGCAAGTGGTCAAAAAG, whereas the sense PCR primer for G2A was AGATTCGCGGCCGCACCATGGCCGGCAAGT-GGTCAAAAAG; the antisense PCR primer was CCGGAGTACTTCAAGAACTGGAATTCTAAGCA. The purified PCR product and pcDNA3.1-Nef_SF2_-V5-VC were digested with *Eco*RI and *Not*I (NEB), and the DNA was isolated by column purification (Zymo Research). The digested pcDNA3.1-Nef_SF2_V5-VC was treated with shrimp alkaline phosphatase (NEB), then ligated with the PCR products overnight at 16°C and transformed into TOP10 competent cells (Thermo Fisher Scientific). Plasmid DNA was isolated from overnight bacterial cultures and verified via Sanger sequencing. For *in vitro* binding studies, HIV-1 NL4-3 Nef (residues 25 to 206) was fused to either SERINC5 intracellular loop 4 (ICL4; residues 332 to 387) or a SERINC5 ICL4 ΔEDTEE mutant using a long, flexible linker. The cDNAs were then cloned into an expression vector pMAT9s (Addgene Plasmid # 112590) using the *NcoI* and *HindIII* sites, fusing SERINC5 Nef-ICL4 to maltose binding protein (MBP-Nef-ICL4). MBP-Nef alone was cloned similarly.

### Expression, purification, and analysis of GST-SERINC3-Loop10, GST-SERINC5-ICL4, MBP-µ1 and MBP-µ2 proteins

We previously reported expression, purification, and analysis of GST-SERINC3-Loop10 (a loop analogous to SERINC5 intracellular loop 4) (47). Here, we expressed, purified, and analyzed GST-SERINC5-ICL4 similarly. SERINC5-ICL4 was cloned into the pGEX4T1 Vector (GE Life Sciences) with an N-terminal GST tag, and the GST-SERINC5-ICL4 construct was transformed into *E.coli* BL21(DE) cells for protein expression. To express phosphorylated GST-SERINC5-ICL4, the construct was co-expressed with both the α and β-subunits of Casein Kinase II (CK-II). The cells were grown to OD600 of ~0.6, induced with 0.1mM IPTG overnight at 16°C, then collected by centrifugation. Cell pellets were lysed using a French press homogenizer. Lysates were clarified by centrifugation at 14,000 RPM. GST-SERINC5-ICL4 was purified using GST-affinity chromatography, HiPrep-Q anion exchange and S200pG chromatography. Purfied GST-SERINC5-ICL4 co-expressed with CK-II was subjected to LC/MS and analyzed as previously reported (47). For the expression of MBP-µ1 and MBP-µ2, we used previously described constructs (48) encoding truncated versions of µ1 (residues 158-423) and µ2 (residues 159-435). These proteins were co-expressed with the pGro7 chaperone in BL21 cells, which was induced by 1.5g/L L-(+)-arabinose at OD600~0.2. To express MBP-µ1 and MBP-µ2, the cells were further grown to OD600~0.6 and induced with 0.1mM IPTG overnight at 16°C. Cells were lysed using a French press homogenizer. The cell lysate was clarified by centrifugation at 14,000 RPM. The proteins were purified using His-Select Nickel affinity gel, Hi Prep-S cation exchange and S200pg gel-filtration chromatography.

### Recombinant Protein Expression for Nef-SERINC5-AP2 pulldowns

For protein expression, *E. coli* BL21(DE3) cells were transformed with the MBP-Nef-ICL4 construct or mutant, grown to OD_600_ ~ 0.8, induced with 0.3 mM IPTG, and expressed overnight at 18°C. Cell pellets were harvested by centrifugation and flash-frozen in liquid nitrogen for storage. GST-tagged μ2_CTD_-truncated AP-2 was prepared as previously described (49): *E.coli* cells overexpressing all four AP-2 subunits were lysed by microfluidization, cell-debris removed by ultracentrifugation, and the supernatant applied to Ni-NTA agarose followed by glutathione-agarose affinity column (GSTrap HP, GE Healthcare). AP-2-containing fractions were pooled, concentrated, and dialyzed into glutathione-free buffer overnight. MBP-Nef and MBP-Nef-ICL4 proteins were purified by Ni-NTA agarose, anion exchange chromatography (HiTrap Q, GE Healthcare), and Superdex 200 size exclusion chromatography.

### GST-pulldowns using recombinant proteins

Purified, GST-SERINC3-Loop10, GST-SERINC5-ICL4, MBP-µ1 and MBP-µ2 proteins were used for *in vitro* GST-pulldowns. An equimolar ratio of GST-tagged proteins were mixed with MBP-µ1 or MBP-µ2, and these mixtures were incubated with GST-resin overnight at 4°C. The next morning, the GST-resins were extensively washed with 20mM Tris-HCL pH 7.5 to remove unbound proteins. The GST-bound proteins were eluted with 10mM glutathione reduced in 50mM Tris HCL pH 8.0. The eluted fractions were analyzed by SDS-PAGE using Coomassie Blue stain. For GST-tagged AP-2 pulldown assays, GST-tagged AP-2 (0.2 mg) and either MBP-tagged Nef or the Nef-ICL4 fusion proteins (0.4 mg, 5-fold molar excess) were mixed in a final volume of 100 μL. Reaction mixtures were loaded onto small, gravity flow columns containing 0.2 mL glutathione Sepharose 4B resin (GE Healthcare), and incubated for 1 hour at 4°C. Protein mixtures and resin were washed extensively with 5x (400 μL) GST binding buffer (50 mM Tris pH 8.0, 100 mM NaCl, 0.1 mM TCEP), and bound protein complexes were eluted with 4x (200 μL) GST elution buffer containing 10 mM reduced glutathione. Elution fractions were analyzed by SDS-PAGE and stained with Coomassie Blue.

### Measurement of Viral Infectivity

Infectivity was measured in virions produced from HEK293 cells or Jurkat TAg *SERINC3/5* KO cells co-transfected with either an infectious molecular clone of HIV-1 (pNL4-3) or a mutant version harboring a deletion in the *nef* gene (pNL4-3ΔNef), and increasing concentrations of pBJ5-SERINC5-iHA or pBJ5-SERINC5-iHA ΔEDTEE. HEK293 cells were seeded at a density of 5×10^5^ cells/ml/well (12-well plates). The cells were transfected the following day with a total of 1.6 µg plasmid, comprising 1.3 µg pNL4-3 or pNL4-3ΔNef, and increasing concentrations of SERINC5-iHA (as indicated), or empty plasmid (pBJ5), using Lipofectamine 2000 transfection reagent, according to the manufacturer’s instructions (Thermo Fisher Scientific). For experiments including glycoGag (Figure 8), HEK293 cells were transfected with a total of 1.6 µg plasmid; 625 ng pNL4-3ΔEnv (Nef+) pNL4-3ΔEnvΔNef (Nef-), 325 ng pVRE (expressing Env), with or without 100 ng pBJ5-HA-gg189 (glycoGag), and increasing concentrations of SERINC5-iHA (as indicated), or empty plasmid (pBJ5). Cells and supernates were harvested after 24 hr. 3.75×10^5^ JTAg S3/5 KO cells in 2.5ml medium were co-transfected (Jurkat-In; MTI Global Stem) with a total of 1.25 µg DNA, comprising 1 µg pNL4-3 or pNL4-3ΔNef, and increasing concentrations of SERINC5-iHA (as indicated), or empty plasmid (pBJ5). Cells and supernates were harvested 48 hr post-transfection. For experiments comparing SERINC5 mutants, a single concentration of pBJ5-SERINC5-iHA, pBJ5-SERINC5-HA ΔEDTEE, pBJ5-SERINC5-iHA AATAA or pBJ5-SERINC5-iHA EDAEE was used (as indicated in the figure legends). Virions were harvested from supernates by centrifugation through a 20% sucrose cushion at 23,500 × g for 1 h at 4°C. The virus pellet was resuspended in culture medium and dilutions used to infect the reporter cell line HeLa P4.R5 in duplicate in a 48-well format. These cells express the HIV-1 co-receptors CD4 and CCR5 and possess a Tat-inducible β-galactosidase gene under the transcriptional control of the HIV-1 LTR. 48 hr post-infection, the cells were fixed with 1% formaldehyde and 0.2% glutaraldehyde for 5 min at room temperature and then stained with 4 mM potassium ferrocyanide, 4 mM potassium ferricyanide, 2 mM MgCl_2_, and 0.4 mg/ml X-Gal (5-bromo-4-chloro-3-indolyl-β-D-galactopyranoside) overnight. Infectious centers (IC) were imaged and quantified using image analysis software (50). The IC data were normalized to the concentration of p24 antigen in each viral stock measured by ELISA (ABL Bioscience). For experiments evaluating the activity of glycoGag, infectivity was measured using HeLa-TZM-bl cells, which contain a luciferase gene under the transcriptional control of the HIV-1 LTR. HeLa-TZM-bl cells were infected with diluted virus stock in duplicate wells of 96-well plates for 48 hours. The culture medium was removed, and the cells were lysed in luciferase reporter gene assay reagent (Britelite, Perkin Elmer); luciferase activity was measured using a luminometer as relative light units (RLU), and normalized to the p24 concentration. To eliminate residual Nef-phenotype in the absence of transfected SERINC5-expression plasmids, the IC/ng p24 or relative light units RLU/ng p24 were expressed relative to the no-transfected-SERINC5 control for each viral genotype, setting the no-transfected-SERINC5 values to 100% for both wildtype and ΔNef in each experiment.

### SERINC5 Virion Incorporation and Western blots

An aliquot of virions purified as described above was used to measure virion-incorporation of SERINC5. The samples of virions were lysed in 30 µl 1x Laemmli Buffer containing 50 mM TCEP (tris(2-carboxyethyl) phosphine; Sigma) and subjected to standard SDS-PAGE after adjustment to equal amounts of p24, as measured by ELISA. Cellular samples from all experiments were lysed in extraction buffer (50mM NaCl, 1% Triton-X-100, 50mM Tris, pH 8.0), nuclei pelleted by centrifugation, and total protein concentration of supernatant was measured by BCA assay (Thermo Fisher Scientific). Equal protein concentrations were mixed with 2x Laemmli buffer containing 100 mM TCEP. To avoid boiling and consequent aggregation of SERINC5 (28, 29), the samples were sonicated (Diagenode Bioruptor) before protein separation by SDS/PAGE and Western blotting. The cell lysates and viral pellets were resolved on 10% denaturing SDS-PAGE gels, transferred onto polyvinylidene difluoride (PVDF) membranes, immunoblotted with the indicated antibodies, and visualized using Western Clarity detection reagent (Bio-Rad). Chemiluminescence was detected using a ChemiDoc Imager System (Bio-Rad). Primary and secondary antibodies were prepared in antibody dilution buffer, consisting of 2% milk in PBST (PBS with 0.02% Tween 20). The following antibodies were used for detection of proteins of interest: HA.11 (mouse, Biolegend), β-actin (mouse, Sigma), GAPDH (mouse, Genetex), HIV-1 p24 (mouse, Millipore) and HIV-1 Nef (sheep) (51).

### Flow Cytometry

Surface SERINC5 was measured in HEK293 cellls and JTAg S3/5 KO cells transfected to express pNL4-3 or pNL4-3ΔNef and pBJ5-SERINC5-iHA or the indicated mutants: ΔEDTEE, EDAEE or AATAA. HEK293 cells were transfected with 1.6 µg total plasmid, 100 ng of which was pBJ5-SERINC5-iHA, and JTAg S3/5 KO cells were transfected with a total of 1.25 µg plasmid, of which 250 ng was pBJ5-SERINC5-iHA. HEK293 cells were stained 24 hours post-transfection and JTAg S3/5 KO cells 48 hours post-transfection. The cells were then washed with ice-cold FACS Buffer (1x PBS + 3% FBS) before staining with mouse anti-HA (diluted 1:200, Biolegend) for 30 minutes on ice. The cells were pelleted by centrifugation and washed in FACS buffer before incubation with goat anti-mouse Alexa Fluor 647 (diluted 1:200, Biolegend) for 30 minutes on ice. To detect intracellular p24, the cells were washed in FACS buffer and fixed and permeabilized with Cytofix/Cytoperm reagent (BD Biosciences) and stained with an anti-p24 FITC antibody (clone KC57; Beckman Coulter) for 30 minutes on ice (diluted 1:100 in Perm wash buffer, BD Biosciences). The cells were washed with FACS buffer and PBS before analysis by flow cytometry. Surface SERINC fluorescence was quantified in at least 1×10^4^ p24-positive cells per condition. For the BiFC assays, HEK293 cells were transfected with 0.8 µg of either a single Venus-N or Venus-C plasmid with an empty vector, or pairwise with Venus N- and C- terminal fusion proteins. Relative fluorescence intensity was measured in 1×10^4^ cells per condition, 24 hours after transfection. Data were collected on a BD Accuri C6 Cytometer and analyzed using C-Flow sampler (BD) and FlowJo (v10, FlowJo LLC) Software.

### Data Analysis and Presentation

Quantitative analyses were performed as described above. Replicate datasets were combined in Microsoft Excel and Graphpad Prism 5.0 software. Figures were produced using Adobe Photoshop and Adobe Illustrator (CS3) software.

## Results

### The EDTEE sequence reduces the sensitivity of SERINC5 to HIV-1 Nef

An alignment between human and various primate SERINC5 proteins revealed a highly conserved acidic sequence (EDTEE; Figure 1) located in a long, predicted cytoplasmic loop (designated intra-cytoplasmic loop 4 (ICL4)). The EDTEE sequence is reminiscent of an acidic cluster membrane trafficking signal, therefore we hypothesized that it might be a Nef-response sequence and support Nef-activity. To test this, we co-transfected HEK293 cells or Jurkat TAg cells that lack endogenous SERINC3 and 5 (*SERINC* 3/5 KO) with pNL4-3 (an HIV-1 infectious molecular clone) or its Nef-negative counterpart (pNL4-3 ΔNef) along with increasing amounts of plasmids expressing either SERINC5-HA (pBJ5-SERINC5-iHA) or SERINC5-HA ΔEDTEE (pBJ5-SERINC5-iHA ΔEDTEE). The virions produced were partially purified by centrifugation through a 20% sucose cushion, and their infectivity was measured using an infectious center (IC) assay. The IC values were divided by the concentration of p24 capsid antigen, and the IC/p24 ratios were normalized to the “no-added SERINC5” control, setting that control value to 100% for both the wild type and the Nef-negative viruses. The latter normalization removed from the presented data differences between the infectivity of wild type and Nef-negative viruses that were not due to the experimental expression of SERINC5 or related mutants by transfection. For our HEK293 cells, the *nef*-infectivity-phenotype in the absence of plasmid-mediated expression of SERINC5 was 6- to10-fold and was presumably due to the endogenous expression of SERINC family members (data not shown). For the Jurkat TAg *SERINC* 3/5 KO cells, the *nef*-infectivity-phenotype in the absence of plasmid-mediated expression of SERINC5 was approximately 2-fold, despite the genetic disruption of both *SERINC* family members with known anti-viral activity (data not shown).

**Figure 1:**
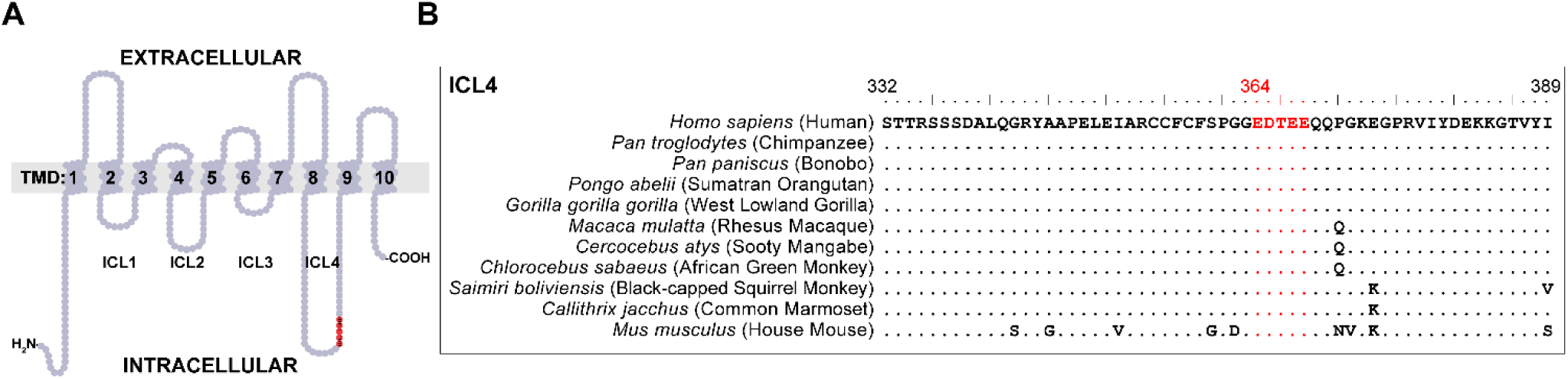
An acidic cluster motif (EDTEE) is highly conserved within human and non-human primate SERINC5. A.) Predicted topology of SERINC5 showing ten transmembrane domains and six cytoplasmic domains, four of which form loops. The EDTEE sequence is shown in red and is found within the long cytoplasmic loop-4 (ICL-4). B.) Amino-acid sequence alignment of SERINC5 ICL-4. The human sequence, the sequences of several non-human primates, and the murine sequence are shown. The conserved EDTEE acidic cluster motif is shown in red.

As expected, we observed a dose dependent antiviral effect of SERINC5, which was greater when virions were produced in the absence of Nef (Figure 2A). Surprisingly, deletion of the EDTEE sequence enhanced the sensitivity of SERINC5 to Nef, but it did not affect the inhibitory activity of the protein in the absence of Nef (Figure 2A). The difference in sensitivity to Nef was not clearly attributable to differences in SERINC5 protein expression within viral producer cells (Figure 2B, which shows a representative experiment in which Jurkat cells were used to produce virions). However, a subtle influence of the deletion on protein-expression was apparent in the dose-response western blot data: the ΔEDTEE mutant seemed slightly underexpressed in the absence of Nef, although it was expressed equivalently to wild type SERINC5 in the presence of Nef. Consistent with the infectivity data, deletion of the EDTEE sequence caused substantially enhanced exclusion of SERINC5 from virions by Nef (Figure 2C, which again shows a representative experiment in which Jurkat cells were used to produce virions). Notably, we confirmed that a 55 kDa form of SERINC5, while the minority species in cells, is the predominant form in virions, an effect due to the selective incorporation into virions of a form of the protein modified by complex glycans (52). Here, the data suggested that the ΔEDTEE mutant might be slightly less efficiently incorporated into virions than wild type SERINC5 in the absence of Nef, although the difference is subtle. Overall, these data support the current model that Nef-mediated exclusion of SERINC5 from virions correlates with enhanced infectivity (28, 29). The data further indicate that the EDTEE sequence within SERINC5 provides a degree of resistance to Nef-activity; Nef is more active in the absence of this sequence.

**Figure 2:**
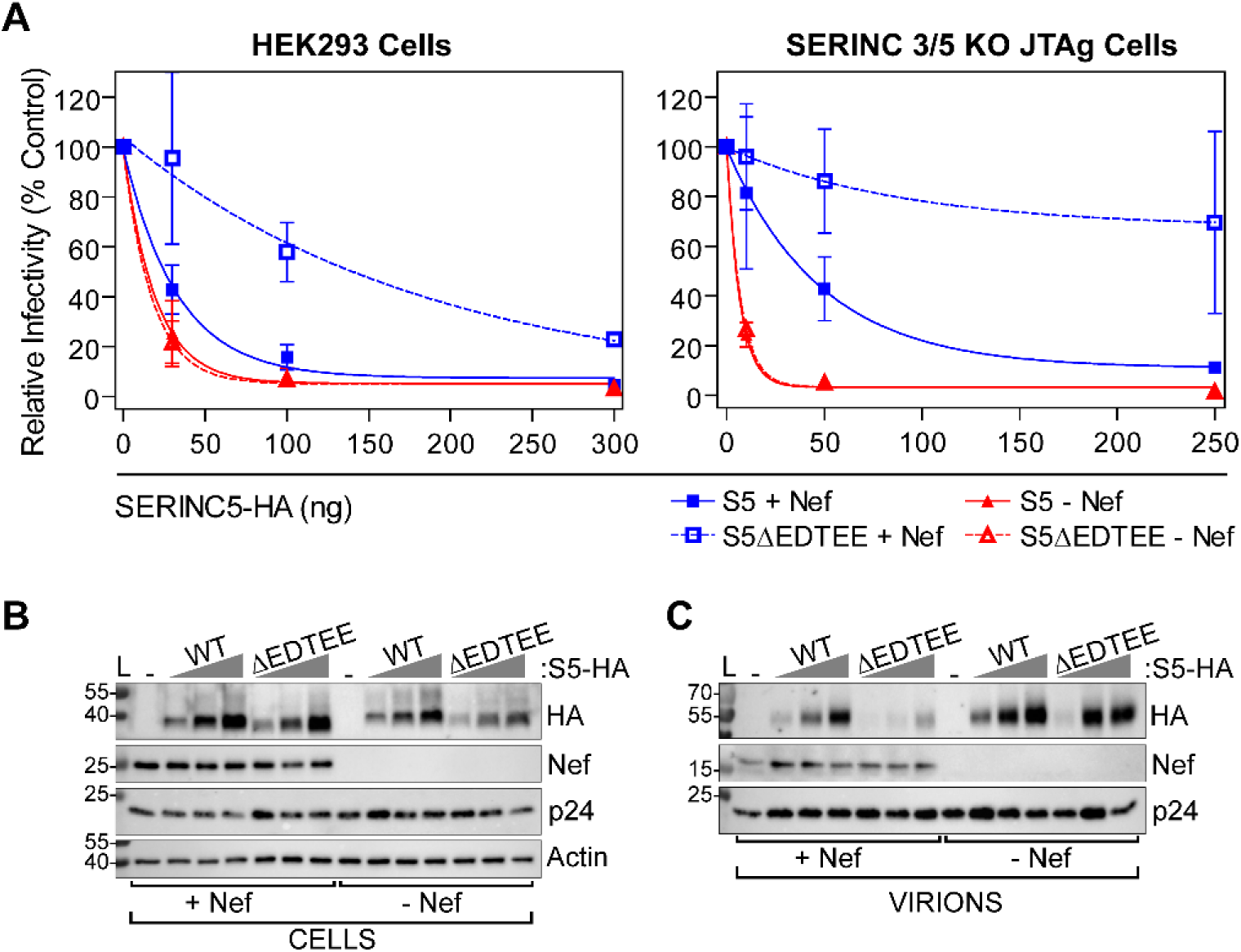
The EDTEE sequence within SERINC5 is not necessary for antiviral activity but confers relative resistance to Nef. A.) HEK293 cells or Jurkat TAg cells lacking SERINC3 and 5 (JTAg *S3/5* KO) were transfected to express NL4-3 (WT) or a *nef*-negative mutant (NL4-3ΔNef) and increasing doses of SERINC5-HA or a mutant lacking the acidic cluster motif (SERINC5-iHAΔEDTEE), as indicated. The produced virions were partially purified by centrifugation through a sucrose cushion and used to infect HeLa P4.R5 cells, which express an LTR-β-galactosidase indicator. Forty-eight hours later, the cells were stained with X-gal, and infectious centers (IC) imaged and quantified. The IC/ml were divided by the concentration (ng/ml) of p24 antigen measured in the virion-preparations by ELISA. The infectious centers per nanogram (IC/ng) were normalized to the no-added-SERINC5 control for each viral genotype (wild type: “+ Nef” or Nef-negative: “- Nef”. Data are presented as the mean percentage relative infectivity, error bars are the standard deviation (s.d.) from *n=2* (HEK293 cells) and *n=3* (JTAg *S3/5* KO cells) experiments. B.) Protein from whole JTAg *S3/5* KO cell lysates from the experiment of panel A were subjected to SDS-PAGE and Western blotting. Membranes were probed with antibodies to detect SERINC5 (HA), Nef, p24, and β-actin. C.) Virions produced by JTAg *S3/5* KO cells and analyzed in the experiment of panel A were normalized by p24 content before SDS-PAGE. Membranes were probed for p24/55, Nef, and SERINC5 (HA). Nef is cleaved within virions by the viral protease, yielding a 16 kDa C-terminal product (51).

### The EDTEE sequence is phosphorylated by casein kinase II in vitro but does not interact with the μ subunit of AP-2

We reported recently that a phosphoserine acidic cluster (PSAC) motif of sequence SGASDEED is present in a cytoplasmic loop of SERINC3 analogous to the loop that contains the EDTEE sequence in SERINC5. Unlike the EDTEE sequence, the SGASDEED sequence has no impact on sensitivity to Nef, despite that the serines of this sequence are under positive selection (38, 47). The SGASDEED sequence of SERINC3 has potential as a membrane sorting or trafficking sequence, however, because it binds the medium (μ) subunits of AP-1 (μ1) and AP-2 (μ2) in a serine-phosphorylation dependent manner (47). Here we observed that when the recombinant SERINC5 loop containing the EDTEE sequence (ICL4) was co-expressed as a GST-fusion protein together with casein kinase II in *E.coli*, the threonine of the EDTEE sequence, as well as upstream serines in the loop, were phosphorylated (Figure 3A). Nonetheless, unlike the analogous loop of SERINC3, phosphorylated SERINC5 ICL4 did not bind to recombinant μ2 *in vitro* (Figure 3B). These data indicate that although the EDTEE sequence has the potential for threonine-phosphorylation, *in vitro* binding data do not support a role as a μ-binding clathrin-adaptor sorting signal.

**Figure 3:**
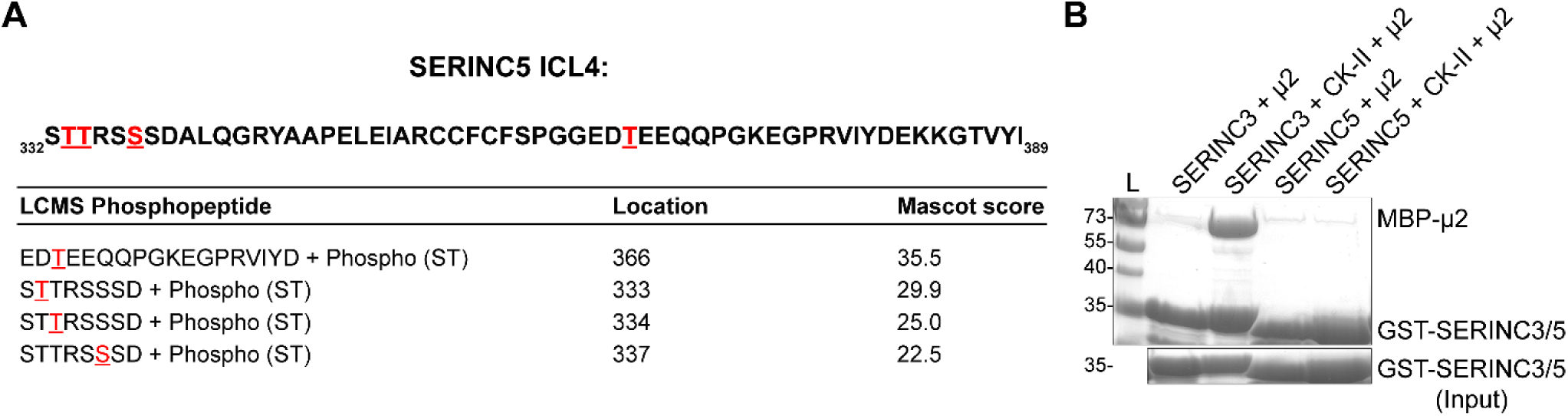
Phosphorylated SERINC5-ICL4 does not bind directly to the medium (µ) subunit of AP-2. A) Sequence of SERINC5-ICL4. GST-SERINC5-ICL4 was co-expressed with casein kinase II (CK-II) in *E.coli*, purified, and analyzed by liquid chromatography/mass spectrometry (LC/MS). Phospho-peptide sequences are shown, and sites of serine-threonine (ST) phosphorylation are highlighted in red and the residue locations are indicated. Mascot scores for the peptide matches are shown. B) Serinc5-ICL4 does not bind to µ2. GST-SERINC3 loop 10 (a positive control) and GST-SERINC5-ICL4 fusion proteins were expressed in *E. coli* either with or without CK-II and tested for binding to µ2 in GST-pulldown assays. The µ2 protein is N-terminally truncated and fused to maltose binding protein (MBP) as a solubility tag. SDS/PAGE gels were stained with Coomassie Blue.

### The acidic residues within the EDTEE motif but not the threonine affect Nef-sensitivity

We next sought to determine whether the acidic nature of the SERINC5 EDTEE motif is the determinant of sensitivity to Nef. We created SERINC5 mutants that either lacked (EDTEE mutated to AATAA) or preserved (EDTEE mutated to EDAEE) acidic residues and tested their restrictive activity and sensitivity to Nef. SERINC5-AATAA, but not SERINC5-EDAEE, was characterized by a relatively enhanced sensitivity to Nef that was similar to the phenotype of SERINC5-ΔEDTEE in both HEK293 and JTAg *S3/5* KO cells (Figure 4A). All the SERINC5 mutants were as restrictive as the wild type protein; that is, they inhibited the infectivity of virions produced in the absence of Nef as effectively as wild type SERINC5, despite that the expression of the ΔEDTEE and AATAA mutants (but not the EDAEE mutant) seemed slightly reduced. Overall, these data suggest that the relative acidity or negative charge of the SERINC5 EDTEE region affects sensitivity to Nef but not intrinsic restrictive activity. The data also indicate that the threonine alone is not a substantial determinant of sensitivity to Nef, even though it would contribute to the negative charge of the region if phosphorylated.

**Figure 4:**
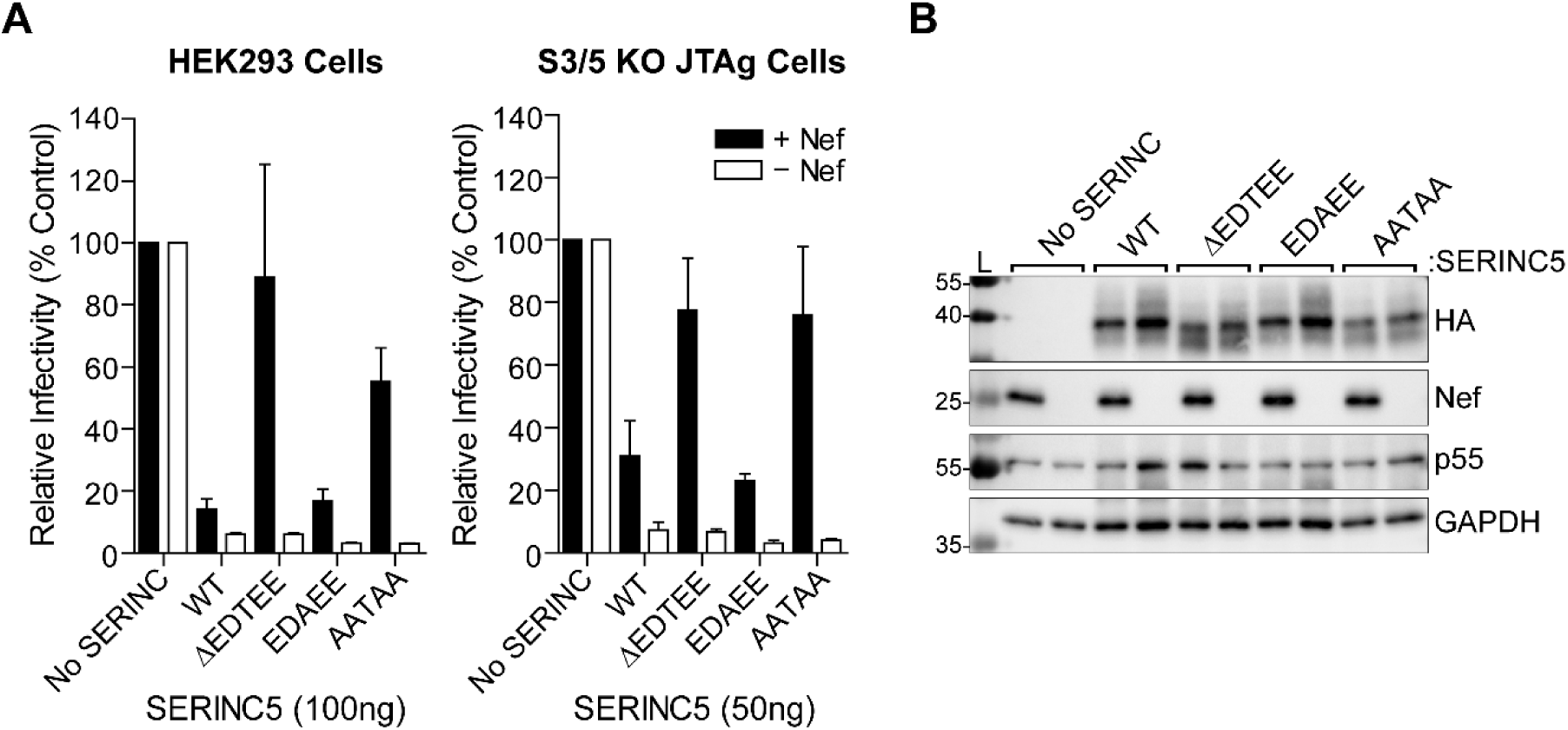
The acidic residues within the SERINC5 EDTEE sequence, but not the threonine, are important for resistance to Nef. A.) HEK293 cells or JTAg *S3/5* KO cells were transfected with pNL4-3 or pNL4-3ΔNef and indicated amounts of plasmid expressing either WT SERINC5 or the following mutants: ΔEDTEE, EDAEE, or AATAA. Virions were harvested and infectivity assays performed in HeLa P4.R5 cells, as described in Figure 2. The infectious centers per nanogram (IC/ng) were normalized to the no-added-SERINC5 control for each viral genotype. Data are presented as the mean percentage relative infectivity, error bars are the s.d. from *n=2* (HEK293 cells) and *n=3* (JTAg *S3/5* KO cells) experiments. B.) Protein derived from whole JTAg *S3/5* KO cell lysates from the experiment of A were subjected to SDS-PAGE and Western blotting. Membranes were probed with antibodies to detect SERINC5 (HA), Nef, p24/55, and GAPDH.

### Deletion of the EDTEE sequence enhances Nef’s ability to downregulate SERINC5

Nef co-opts endocytic machinery, namely AP-2 and clathrin, to downregulate SERINC5 from the plasma membrane (28, 29). We reasoned based on our virologic data that the ability of Nef to downregulate SERINC5 would be increased by deletion of the EDTEE sequence. To study the downregulation of cell surface SERINC5 by Nef, we co-transfected cells with pNL43 or pNL43ΔNef together with the SERINC5-iHA expression-plasmids and measured surface SERINC5 levels by immunofluorescent staining and flow cytometry. All of the SERINC5 constructs (wild type, ΔEDTEE, AATAA, and EDAEE) were similarly expressed at the cell surface in Jurkat TAg *SERINC*3*/5* KO cells in the absence of Nef (Figure 5A, lower panel). SERINC5-ΔEDTEE and SERINC5-AATAA, but not SERINC5-EDAEE, were downregulated more efficienty than wild type SERINC5 in both JTAg *S3/5* KO and HEK293 cells (Figure 5). These results are consistent with the virologic data and support the correlation between the downregulation of cell surface SERINC5 and the enhancement of infectivity by Nef. The data further support that the EDTEE sequence provides relative resistance to Nef-mediated modulation of SERINC5.

**Figure 5:**
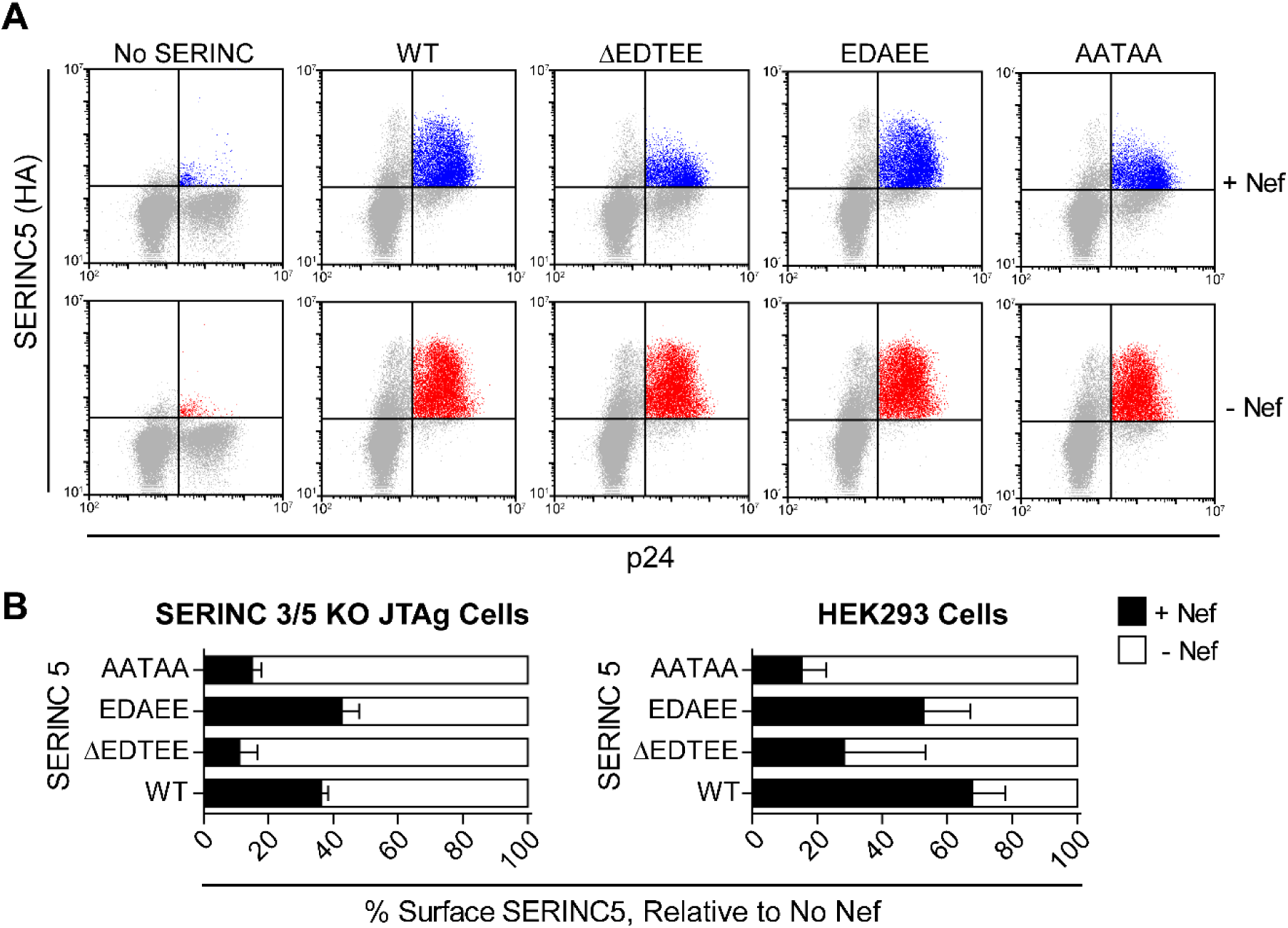
Mutation of the EDTEE sequence enhances the downregulation of SERINC5 from the cell surface by Nef. JTAg *S3/5* KO or HEK293 cells were transfected with pNL4-3 (+ Nef) or pNL4-3ΔNef (- Nef) and either 250 ng (JTAg *S3/5* KO) or 100ng (HEK293) of plasmid expressing WT SERINC5 or the EDTEE mutants: ΔEDTEE, EDAEE, or AATAA. The cells were stained for surface SERINC5 (HA, Alexa Fluor-647) and intracellular p24 (FITC). A.) Representative two-color dot-plots showing surface expression of SERINC5 (HA) in p24-positive JTAg *S3/5* KO cells (+/- Nef). B.) The MFI of surface SERINC5 (HA) in p24-positive cells was quantified +/- Nef. Data are presented as mean percentage MFI normalized to no-Nef control, error bars are the s.d. of *n=2* (JTAg *S3/5* KO cells) or *n=4* (HEK293 cells) experiments.

### Deletion of the SERINC5 EDTEE sequence does not enhance interaction with Nef

We used a recently reported bimolecular fluorescence complementation (BiFC) assay (34, 46) to test the hypothesis that the interaction of Nef and SERINC5 is enhanced in the absence of the EDTEE sequence. In this assay, the two proteins of interest are fused to either the N- or C-terminus of Venus (yellow fluoresecent protein). A fluorescent signal is generated if the two proteins interact, enabling the quantitative measurement of protein-protein interactions within living cells. Here, we fused the N-terminus of Venus to the C-terminus of either wild-type NL4-3 Nef or a myristolyation-signal mutant incapable of associating with membranes (Nef G2A) (6). We also fused the C-terminus of Venus to the C-terminus of either SERINC5, SERINC5ΔEDTEE, SERINC5 EDAEE or SERINC5 AATAA. These constructs were used to transfect HEK293 cells either singly or in pairs, and the relative fluorescence was measured by flow cytometry twenty-four hours later. We detected modest fluorescence when only Nef-VN or SERINC5-VC was expressed and a 4-fold relative increase in fluorescence when these two proteins were co-expressed (Figure 6A). This increase of the fluorescent signal was lost when SERINC5-VC was paired with Nef-G2A-VN, consistent with the notion that Nef requires membrane-association to interact with SERINC5. We did not detect an increased interaction-signal when Nef-VN was paired with either SERINC5-ΔEDTEE-VC or SERINC5-AATAA-VC relative to SERINC5-VC (Figure 6A). No differences in the expression of these fusion proteins was detected by western blot (Figure 6B). These data suggest that deletion of the SERINC5 EDTEE sequence does not enhance its interaction with Nef.

**Figure 6:**
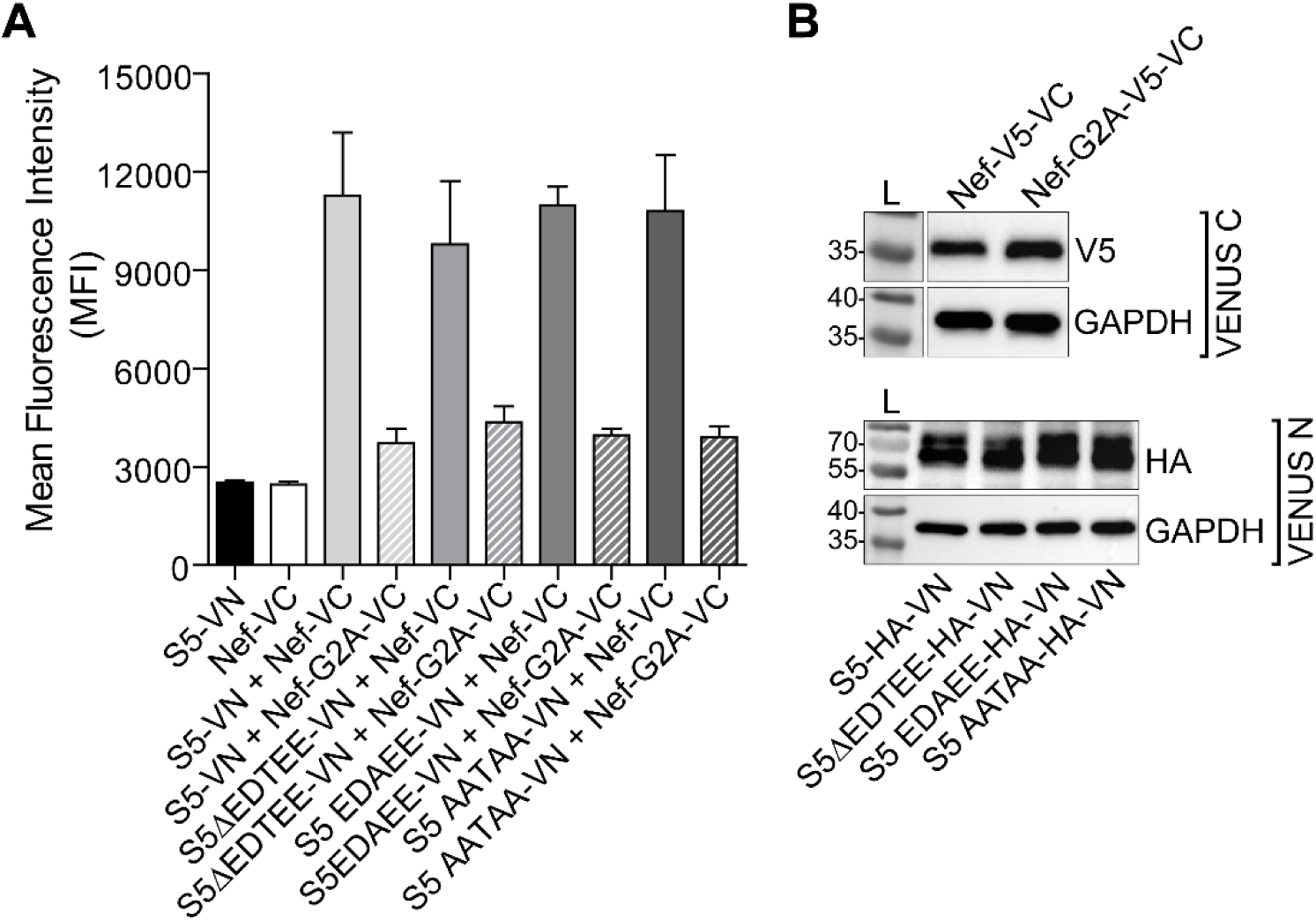
Intracellular interaction between Nef and SERINC5 as measured by bimolecular fluorescence complementation (BiFC) is not substantially affected by the EDTEE sequence. A.) HEK293 cells were transfected with plasmid constructs expressing Nef-Venus-C (VC), Nef G2A-VC, SERINC5-Venus-N (VN), SERINC5ΔEDTEE-VN, SERINC5 EDAEE-VN or SERINC5 AATAA-VN either singly or in the pairs indicated. Twenty-four hours later, the fluorescence intensity (FL1) was measured by flow cytometry. The data are presented as the mean fluorescence intensity of Venus signal, error bars are s.d. for *n=3* independent experiments. B.) Protein from whole cell lysates from a representative experiment was subjected to SDS-PAGE and Western Blotting. Membranes were probed with anti-V5 (Nef-VC), HA (SERINC5-VN) or GAPDH (loading control) antibodies.

### Deletion of the EDTEE sequence does not enhance binding of a Nef-SERINC5 cytoplasmic loop fusion protein to AP-2 in vitro

We next sought to determine whether deletion of the EDTEE sequence enhances formation of a ternary complex including Nef, a cytoplasmic loop of SERINC5 and AP-2. As noted above, a previous study showed that the long cytoplasmic loop within SERINC5 – ICL4 – confers Nef-responsiveness (39). Because ICL4 contains the EDTEE sequence, we produced recombinant proteins containing NL4-3 Nef (residues 25-206) fused via a long flexible linker to either SERINC5 ICL4 (residues 332-387) or a SERINC5 ICL4 ΔEDTEE mutant (Figure 7A). A protein containing only the Nef component served as a control. Each of these proteins was fused to maltose-binding-protein (MBP) to enhance their solubility. Binding of these proteins to a recombinant, μ2_CTD_-truncated AP-2 heterotetramer *in vitro* was analyzed by pulldown assays using GST-tagged AP-2 mixed wth either MBP-Nef, MBP-Nef-SERINC5 ICL4 or MBP-Nef SERINC5 ICL4 ΔEDTEE (Figure 7B). ICL4 strikingly stimulated the pulldown of Nef with the μ2_CTD_-truncated AP-2 complex, a result consistent with the notion that this cytoplasmic loop is a Nef-response sequence and that Nef and ICL4 together bind efficiently to AP-2. However, we detected little or no influence of the EDTEE sequence in this assay: the Nef-ICL4 fusion protein did not clearly bind more efficiently to AP-2 when the EDTEE sequence was deleted (Figure 7B). These results suggest that the the EDTEE sequence does not interfere with formation of a Nef, SERINC5-ICL4, AP-2 complex when assessed using recombinant proteins *in vitro*.

**Figure 7:**
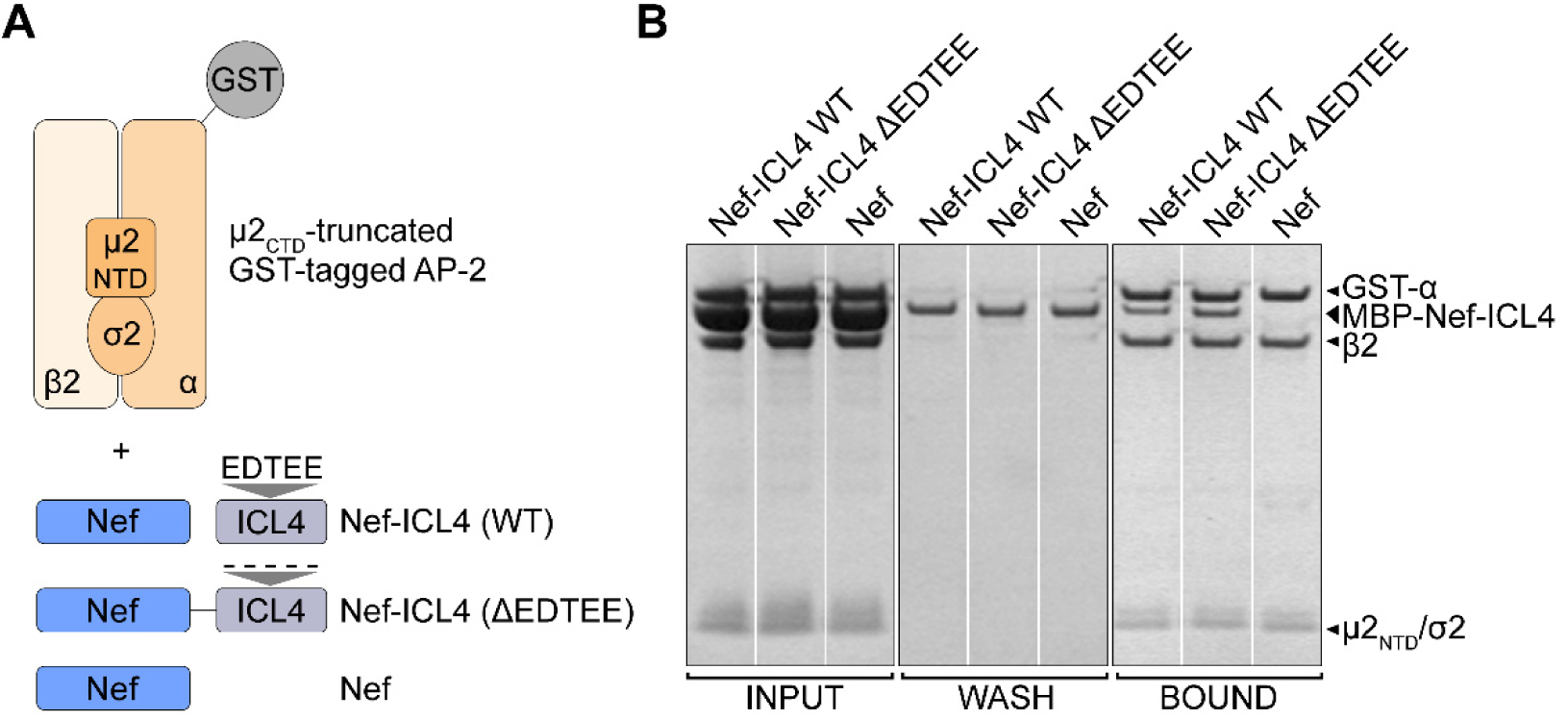
SERINC5 (ICL4) stimulates the interaction of Nef with AP-2 *in vitro* independently of the EDTEE sequence. A.) Schematic of recombinant protein constructs used to study formation of a Nef-SERINC5 ICL4 complex *in vitro*. A heterotetrameric AP-2 complex core with a GST tag on the α subunit; the C-terminal two-thirds of the µ2 subunit is deleted, as are the appendage domains of the α and β2 subunits. Nef was fused to SERINC5 ICL4 (with or without the EDTEE sequence) via a linker peptide; these proteins were further fused to the maltose-binding-protein (MBP) as a solubility tag (not shown). MBP-Nef alone was used as a control. B.) GST-pulldown assay assessing the binding of Nef-SERINC ICL4 (with or without the EDTEE sequence) or Nef alone to the truncated AP2 core *in vitro*. The input protein mixtures, the protein(s) washed through the GST-matrix, and the protein(s) that remained bound to the GST-matrix were run on an SDS/PAGE gel and stained with Coomassie Blue.

### Role of the EDTEE sequence in the antagonism of SERINC5 by glycoGag

Based on our results with Nef, we hypothesized that the EDTEE sequence might affect the antagonism of SERINC5 by the glycosylated Gag (glycoGag) protein of Moloney murine leukemia virus (M-MLV). MLV glycoGag counteracts SERINC3 and 5 and rescues infectivity of *nef*-deficient HIV-1 (35). The majority of the extracellular domain of M-MLV glycoGag is dispensable for this activity (53). We therefore used a minimal active truncated form of glycoGag which contains the N-terminal 189 residues (gg189) to test the ability of glycoGag to rescue the infectivity of HIV-1 lacking Nef in the presence of either SERINC5 or the SERINC5-mutant lacking the EDTEE sequence. For these experiments, HIV-1 Env was provided in *trans* in the virion-producer cells, in order to abrogate syncytia-formation in the target cells, which was strikingly exaggerated when the absence of Nef was complimented by glycoGag and Env was encoded in the viral genome (data not shown). HeLa-TZM-bl indicator cells were used for luminometric measurement of infectivity, as this provided a more sensitive method for measuring the infectivity of the pseudo-virions. We confirmed that, as shown above, the EDTEE sequence provided relative resistance to Nef when infectivity was measured using this modified assay design (Figure 8). Moreover, as reported previously, glycoGag efficiently antagonized the activity of SERINC5 as an inhibitor of infectivity (Figure 8). Unlike Nef, however, the activity of glycoGag was not enhanced when the EDTEE sequence of SERINC5 was deleted. In contrast, the activity of glycoGag against the EDTEE-mutant was slightly diminished. These data suggest that the role of the EDTEE sequence in SERINC5 is Nef-specific, despite that the cellular cofactors involved in SERINC antagonism by Nef and glycoGag appear to be similar (28, 29, 53).

**Figure 8:**
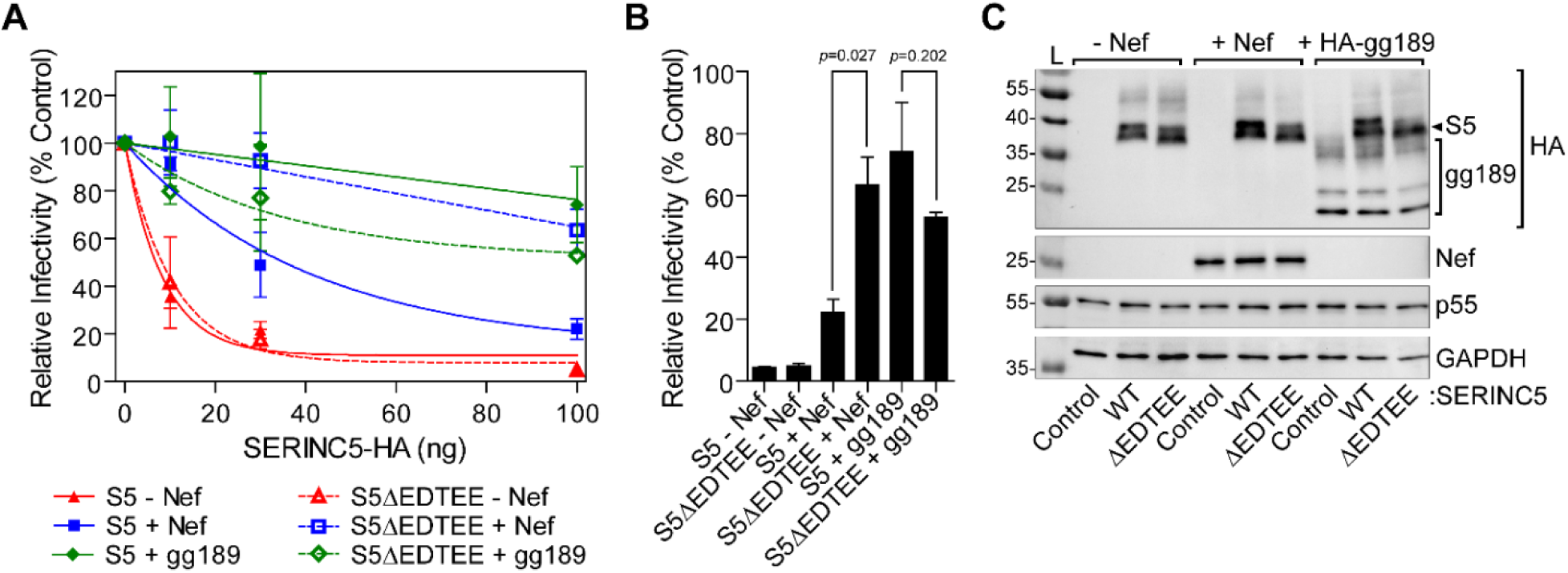
Deletion of the EDTEE sequence does not enhance the activity of MLV Glycogag as an antagonist of SERINC5. A.) HEK293 cells were co-transfected with plasmid constructs encoding indicated amounts of SERINC5-iHA and either *env-* and *nef-*deficient (Nef -), or *env-* deficient (Nef +) HIV-1 provirus. Env expression was provided in *trans* by an NL4-3-derived construct (pVRE). An HA-tagged minimal active glycoGag (HA-gg-189) was also provided in *trans*. Virions were harvested twenty-four hours post-transfection, partially purified by centrifugation through a sucrose cushion, and used to infect HeLa-TZM-bl luciferase indicator cells. Luciferase activity was measured 48 hr post-infection and normalized to p24 antigen (relative light units, RLU/ng p24). Data are expressed as percentage infectivity relative to the no added SERINC control for each condition (with or without Nef or glycoGag). Error bars indicate standard deviation for *n=2* independent experiments. (B) Comparison of relative infectivity of virions in absence or presence of 100 ng SERINC5-iHA (*p*-values derived from Student’s *t*-test are indicated). (C) Protein from whole cell lysates was subject to SDS-PAGE and Western blotting. Membranes were probed with antibodies to detect SERINC5 (HA), glycoGag (gg189, HA), Nef, p55 (Gag), and GAPDH.

## Discussion

We intially hypothesized that the acidic cluster within the long cytoplasmic loop of SERINC5 – the sequence EDTEE in ICL4 – might function as a protein sorting motif in concert with HIV-1 Nef, and thus support Nef-activity as a SERINC5-antagonist. Instead, our data indicate that Nef is more effective as a SERINC5-antagonist in the absence of the EDTEE sequence. A SERINC5 mutant lacking this sequence, or a mutant in which the acidic residues are replaced with alanines (but not a mutant in threonine is replaced with alanine), is more effectively antagonized by Nef at the levels of counteraction of SERINC5-mediated inhibition of infectivity, down-regulation of SERINC5 from the cell surface, and exclusion of SERINC5 from virions. In general, these data are consistent with the current model of surface down-regulation and virion-exclusion of SERINC5 as the basis for Nef-mediated enhancement of infectivity, and they support the notion that the region of SERINC5 containing the EDTEE sequence determines Nef-sensitivity.

While our work was in progress, Dai and colleagues mapped the intracellular cytoplasmic loop that contains the EDTEE sequence (designated ICL4 in their study) as the key region of the protein required for sensitivity to Nef (39). These investigators identified two hydrophobic residues in the N-terminal half of the loop that are required for response to Nef. While our data are consistent with the conclusion that ICL4 contains determinants of Nef-sensitivity, they reveal that in addition to residues that support Nef-activity, the loop also contains a sequence – EDTEE- that is inhibitory to Nef-activity.

Why would a cytoplasmic loop of SERINC5 contain such an inhibitory sequence? Although *SERINC5* does not seem to be under positive selection among primates, a genetic signature of host-pathogen conflict (38), we considered that the EDTEE sequence might have evolved to provide protection against diverse retroviruses and the SERINC antagonists that they encode. The retroviral accessory proteins Nef (found in HIVs and SIVs), glycoGag (found in MLV) and S2 (found in EIAV) are structurally unrelated proteins that all enhance viral infectivity by counteracting SERINC5 (28, 29, 35, 36). However, our data indicate that the EDTEE sequence is not inhibitory to the activity of glycoGag, suggesting that the impact of this sequence is potentially Nef-specific. This scenario weighs against the notion that the protein acquired the EDTEE sequence as a general defense against retroviral antagonists. It also implies that Nef has not yet optimally evolved to counteract SERINC5. Alternatively or in addition, the importance of other Nef-functions for viral fitness might preclude such evolution.

How does the EDTEE sequence affect Nef-responsiveness? One possible explanation is that the sequence is a membrane trafficking signal that directs SERINC5 away from Nef. However, we found no evidence that deletion of the EDTEE sequence influences the subcellular localization of SERINC5 (data not shown), nor does the loop containing the EDTEE sequence bind the µ subunit of the clathrin adaptor AP-2 *in vitro*, as the analogous loop of SERINC3 does.

Another possibility is that the negative charge of the EDTEE sequence inhibits the interaction with Nef. This model is consistent with the requirement of the acidic residues for this phenotype; it might also be consistent with the presence of an acidic cluster in the N-terminal region of Nef, which might repel SERINC5. However, our measurement of the SERINC5/Nef interaction using bi-molecular fluorescence complementation did not support this model: the interaction was unaffected by deletion of the EDTEE sequence. Moreover, a charge-repulsion model predicts that a Nef-mutant in which the acidic cluster is neutralized would be more active as an antagonist of wild type (EDTEE-motif containing) SERINC5, but we did not find that to be the case (data not shown).

Yet another possibility is that a subtle decrease in the steady-state expression of the ΔEDTEE and AATAA mutants is sufficient to increase their apparent Nef-responsiveness. Several lines of evidence weigh against this possibility. First, the SERINC5 mutants appear to reach the plasma membrane, the presumed site of Nef counteraction via endocytosis, as efficiently as the wild type protein (as measured by flow cytometry). Second, none of our data suggest that the intrinsic restrictive activities of the mutant proteins are decreased, weighing against the functional significance of the subtle differences detected in some or our western blots. Third, and perhaps most importantly, the ΔEDTEE mutant was not more responsive than the wild type SERINC5 to glycoGag; rather, the mutant appeared slightly less responsive. The observation that the effect of deleting the EDTEE sequence is opposite when testing responsiveness to Nef versus glycoGag is inconsistent with the notion that the observed virologic phenotypes are consequences of the levels of protein expression.

These considerations leave open the question of exactly how the EDTEE motif in SERINC5 specifically inhibits the activity of Nef as a SERINC5-antagonist. Although our binding experiments using recombinant proteins *in vitro* do not clearly support the hypothesis that the EDTEE sequence inhibits the interaction between Nef-SERINC5 and the AP-2 clathrin adaptor complex, we have not yet attempted to assess this ternary interaction in the more complex environment of human cells. A structural explanation of the interaction between Nef, SERINC5 ICL4, and AP-2 might yet provide an answer for the currently enigmatic role of this motif.

## Acknowledgements

The authors thank Dr. Massimo Pizzato for the pBJ5 and pBJ5-HA-gg189 plasmids; Dr. Heinrich Gottlinger for pBJ5-SERINC5-iHA plasmid and the *SERINC3/5* KO Jurkat TAg cell line; Dr. Yonghui Zheng for the plasmid pcDNA3.1-Serinc5-VN-HA; and Drs. Thomas Smithgall and Sherry Shu for the plasmid pcDNA3.1-Nef_SF2_-V5-VC. We thank The Pendleton Charitable Trust for equipment. P.W.R. was supported by NIH grant K12GM068524. This work was supported by NIH grant R01AI129706 to J.G.

